# Evolutionary capacitance emerges spontaneously during adaptation to environmental changes

**DOI:** 10.1101/101055

**Authors:** Paul G Nelson, Joanna Masel

**Author notes:** The code used to generate all data in this manuscript is available at: https://github.com/pgnelson/emergent_capacitance. Paul G Nelson Department of Ecology & Evolutionary Biology, University of Arizona, Joanna Masel Department of Ecology & Evolutionary Biology, University of Arizona University of Arizona.

## Abstract

All biological populations are to a greater or lesser degree evolvable, but the forces that shape evolvability, especially the evolution of evolvability as an adaptive response to a changing environment, have been a source of controversy. One source of enhanced evolvability is the benign status of “cryptic sequences” typically expressed at low levels due to molecular errors, but with the potential to be expressed more fully following mutational co-option. A genome enriched for benign cryptic sequences has a more benign mutational neighborhood, via the possibility of co-option, and thus enhanced evolvability. Whether selection for evolvability itself can be the cause of a more benign mutational neighborhood remains an open question. Here, we show that environmental change can cause the evolution of increased evolvability, despite our use of a strong-selection weak mutation regime that precludes, by design, the adaptive evolution of evolvability. Instead, enhanced evolvability arises as a byproduct of environmental change via a novel mechanism that we call “emergent evolutionary capacitance”. When the environment changes, increased molecular error rates evolve as a strategy to rapidly change phenotypes, with the side effect of purging deleterious cryptic sequences and enhancing the mutational neighborhood for future adaptation. The behavior is strikingly similar to that seen in a model system for capacitance, the yeast prion [PSI+].

## INTRODUCTION

### Evolvability

Understanding the determinants of a species’ capacity to evolve is a central goal of evolutionary biology. A species’ evolvability, i.e. its ability to generate non-deleterious heritable phenotypic variation, affects its ability to adapt, propensity to speciate, and ultimately its risk of extinction (Wagner and Altenberg 1996; Wagner 2005a; Hendrikse *et al.* 2007). The possibility of adaptive evolution of evolvability (i.e. selection favoring alleles due to their adaptation-enhancing properties) has been posited (Kirschner and Gerhart 1998; Woods *et al.* 2011) and its plausibility has even been shown mathematically (Eshel 1973; Masel and Bergman 2003; King and Masel 2007; Draghi and Wager 2008), but the topic remains controversial (Ruden *et al.* 2003; Sniegowski and Murphy 2006; Lynch 2007b).

A variety of distinct mechanisms can modify or enhance evolvability, including sexual recombination, modularity, robustness, and mutation rate. Sex can increase genetic variation by recombining alleles from different individuals to produce novel and potentially more fit genotypes (Otto and Lenormand 2002). Gene network modularity can establish genetic correlations that avoid deleterious trait combinations (Hansen 2003). Genetic robustness enhances evolvability in two ways. First, more robust populations are predisposed to maintain more conditionally neutral standing variation, providing more potentially adaptive genetic variation should the environment change, either immediately (Waddington 1957; Le Rouzic and Carlborg 2008) or following additional mutations (Wagner 2005b). Second, incomplete robustness to developmental errors causes future mutations to be pre-screened (Masel and Trotter 2010), resulting in a more benign mutational neighborhood (Whitehead *et al.* 2008; Rajon and Masel 2013) or recombinational neighborhood (Masel 2006), and thus enhancing evolvability.

### Evolution of Evolvability

While the impact of these various mechanisms on evolvability is well established, the evolutionary drivers of these evolvability-affecting mechanisms are more controversial. The evolution of mutation rates provides a classic and illustrative example. Mutations are the ultimate source of genetic variation, making mutation rate an important determinant of evolvability (André and Godelle 2006). However, due to a deleterious mutation bias (Eyre-Walker and Keightley 2007), high mutation rates inherently cause a net decrease in offspring fitness. Therefore, all else being equal, the optimal mutation rate is typically zero. Explanations for non-zero mutation rates fall into three general camps: mutations rates evolve as an intrinsic property, as an adaptive consequence of selection for higher evolvability, or as a byproduct of other traits under selection (De Visser *et al.* 2003; Masel and Promislow 2016).

To understand how non-zero mutation rates, and thereby evolvability, evolves as an intrinsic property (Figure 1A), note that the mutation rate is, in part, a function of DNA replication fidelity, which is constrained by the accuracy of DNA polymerases (Fijalkowska *et al.* 2012). Organisms can decrease the mutation rate by utilizing a variety of DNA polymerases to proofread (Fijalkowska *et al.* 2012). However, if the mutation rate is small, any further decrease will have small fitness consequences and may be swamped by mutation bias toward lower fidelity polymerases (Lynch 2012), especially in small populations in which selection for high fidelity is weak. Therefore, evolvability due to a non-zero mutation rate is an intrinsic property of any population subject to mutation bias toward lower DNA replication fidelity. Additionally, as mutation bias tends to lead to higher mutation rates, it is also possible that the extreme population bottlenecks imposed by most natural selection experiments could have led to higher mutation rates simply due to a reduced ability to purge them (Sung *et al.* 2012).

**Figure 1:**
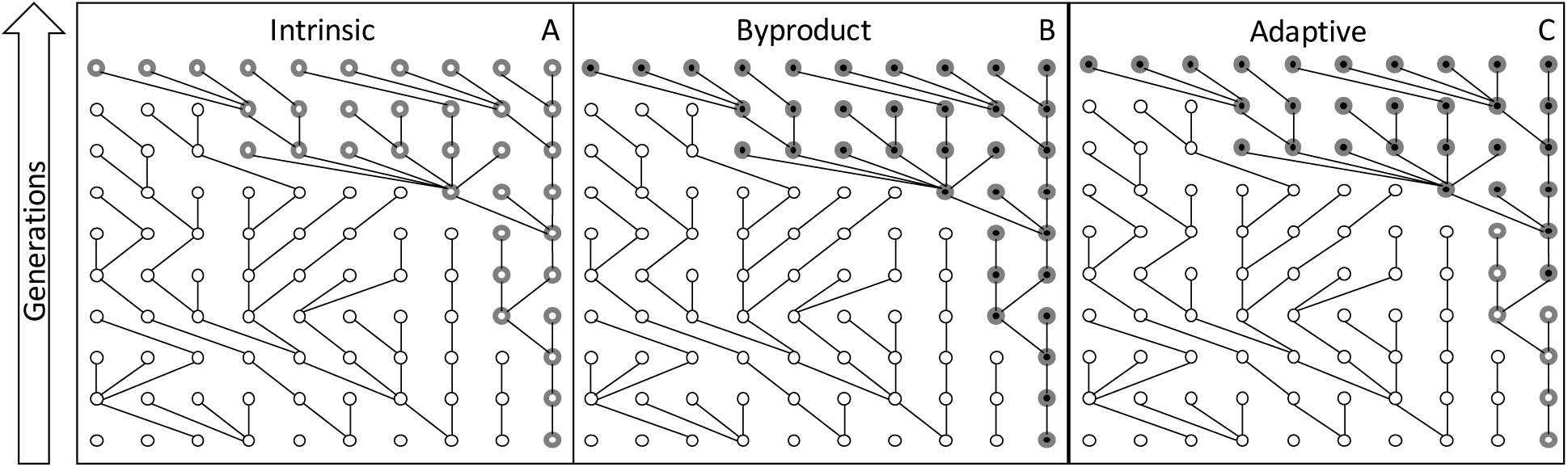
Three scenarios for the evolution of evolvability. Each circle represents an individual in an asexual population of constant size. Initially, the population is composed of nine individuals of the non-evolvable genotype (plain open circles) and one individual possessing an allele that bestows the potential for beneficial mutations (circles with thick grey outline). In panel A, the evolvability allele rises to fixation through drift. In panel B, evolvability is a pleiotropic side effect of an allele that directly increases fitness (filled circle with thick grey outline indicating evolvability), allowing evolvability to fix as a byproduct of its directly adaptive attributes. Finally, C shows the adaptive evolution of evolvability. The evolvability allele bestows no fitness advantage directly, therefore the allele must persist due to drift long enough for the directly beneficial allele to arise (filled circles) and then hitchhike with the directly adaptive allele to fixation.

Alternatively, evolvability due to high mutation rates can arise as a byproduct of other traits under selection (Figure 1B). Given sufficient selection, error rates, including errors in DNA replication, can be made arbitrarily low via kinetic proofreading (Hopfield 1974). However, kinetic proofreading comes with inherent energetic costs; lower error rates entail increasing energetic investment and slower growth. Therefore, selection for rapid growth may result in the evolution of higher mutation rates, and therefore increased evolvability as a byproduct (André and Godelle 2006).

Assertions about the adaptive evolution of evolvability, also known as second order selection (Tenaillon *et al.* 2001; Woods *et al.* 2011), have been especially contentious (Lynch 2007b; Pigliucci 2008). Adaptive evolution of evolvability is criticized as requiring group selection, and because group selection tends to be weak, the adaptive evolution of evolvability is said to be unlikely (Lynch 2007b). However, group selection is a special case of a more universal form of selection, namely lineage selection (Akcay and Van Cleve 2016; Lehmann *et al.* 2016). A lineage begins with a new mutation, and includes all individuals who share that mutation via identity by descent. Lineage selection, i.e. competition between lineages, is therefore, by definition, the determinant of evolutionary outcomes. Group selection is a special case of lineage selection in which members of the same lineage tend to assort with one another in identifiable “groups”. When alleles have only direct effects on fitness, and have the same effect on each individual that bears them, evolutionary outcomes are also well described by traditional measures of individual-based, single-generation, selection coefficients. However, when a trait in one individual affects the success of others in its lineage, either contemporaneously with respect to altruistic behavior, or with a time delay by facilitating the appearance of new adaptive phenotypes, the success of the lineage as a whole must be taken into account (Eshel 1973; Masel and Bergman 2003; Kussell *et al.* 2005; Griswold and Masel 2009). This sometimes leads to cases where lineage selection implies the opposite outcome to that predicted by individual-based selection, including the evolution of altruism (Akcay and Van Cleve 2016; Lehmann *et al.* 2016) and bet-hedging (Philippi and Seger 1989; King and Masel 2007). The concept of lineage selection goes by other names in particular contexts, including clone selection in asexual populations (Eshel 1973), and invasion fitness in a number of contexts, including adaptive dynamics, when the focus is on the rate of short-term increase from rarity (Metz *et al.* 1992; Champagnat *et al.* 2006; Lehmann *et al.* 2016). In a well-mixed and deterministic population, lineage fitness is given by the dominant Lyapunov exponent of a matrix describing changes in both the population and the environment (Metz *et al.* 1992; Kussell *et al.* 2005).

Continuing our example of the evolution of mutation rates, high mutation rates are approximately neutral to an individual. However, high mutation rates increase the likelihood of a highly beneficial mutation later occurring in the mutator lineage. This highly beneficial mutation will, over multiple generations, increase lineage size and may cause the mutator lineage to fix. Thus, when determining the fate a mutator allele, one must take into account the not only the direct effects of the allele, but also the way in which it might, in the future, change the fitness of the genetic background that the allele is likely to be in. For an allele bestowing high mutation rates to be favored by selection, three events must occur. First, a mutator allele arises in the population (Figure 1C, generation one). This high mutation rate allele is not directly adaptive, and must therefore persist due to drift. Then a second, directly adaptive allele arises in an individual with the mutator allele (Figure 1C, generation five). Finally, linkage disequilibrium between the adaptive allele and mutator allele causes the mutation rate allele to sweep to fixation (Figure 1C, generation ten). While the mutator allele does not intrinsically increase fitness in the individual possessing it, it does increase the overall fitness of the lineage through increased evolvability.

Note that fixation of an evolvability-enhancing allele due to drift (Figure 1A), or as a byproduct (Figure 1B), can occur in a scenario with only one polymorphic locus. This is not true for the adaptive scenario; the directly adaptive allele must appear and sweep while the evolvability locus is still polymorphic, so two simultaneous polymorphisms are required for the adaptive evolution of evolvability. Thus, by conducting evolutionary simulations in which the possibility of multiple simultaneous polymorphisms is prohibited by design, the adaptive evolution of evolvability is made impossible, while still allowing enhanced evolvability to arise as a byproduct of other evolutionary dynamics. We achieve this prohibition by using the strong-selection weak-mutation assumption (also known as origin-fixation dynamics), in which each mutation either fixes or goes extinct before the next appears, prohibiting multiple simultaneous polymorphisms (Gillespie 1982; McCandlish and Stoltzfus 2014).

### Evolution of evolvability-promoting high error rates

An ideal system for studying the evolution of evolvability would exhibit clear, preferably binary differentiation between high evolvability and low evolvability states, while still retaining a modicum of biological relevance. To achieve this, we use a model of the co-evolution of cryptic sequences and stop-codon read-through rates, subject to an evolutionary positive feedback loop that yields two distinct outcomes (Rajon and Masel 2011). The positive feedback loop works as follows. When most cryptic sequences are deleterious, selection favors very low read-through rates to avoid their expression. Because read-through rates are low, cryptic sequences are shielded from selection, and under such weak selective conditions, mutation bias keeps them deleterious (Rajon and Masel 2011). Conversely, when most cryptic sequences are benign, selection favors high read-through rates to avoid the costs associated with proofreading; this quickly purges any remaining deleterious mutations in cryptic sequences. This positive feedback loop produces two attractors: mostly deleterious cryptic sequences coupled with low read-through rates, and mostly benign cryptic sequences coupled with high read-through rates. Importantly, the two attractors also differ in their evolvability. A population with mostly deleterious cryptic sequences is limited to adaptation through mutations in the non-cryptic, open reading frame of genes. In contrast, populations with mostly benign cryptic sequences have three potential sources of variation: mutations in regular coding sequences, mutational co-option of cryptic sequences through loss of stop-codons, and *trans* changes in the stop-codon read-through rate (Figure 2). In other words, populations with mostly benign cryptic sequences are more robust to expressing cryptic sequences and therefore have more mutational options, making them more evolvable and better able to adapt to changes in the environment (Rajon and Masel 2011; Rajon and Masel 2013).

**Figure 2:**
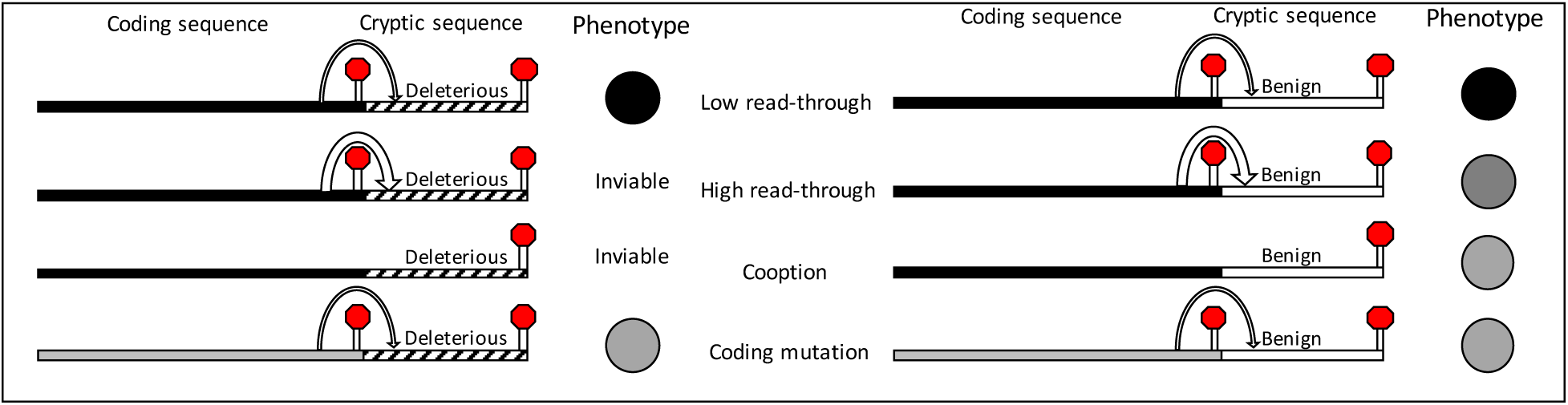
Stop-codon read-through, cryptic sequences, and evolvability. Organisms expressing deleterious cryptic sequences are inviable, leaving mutations in the coding sequence as the only means of adaptation. Benign cryptic sequences, however, can yield potentially useful new phenotypes when expressed. Therefore, organisms with benign cryptic sequences can change phenotype by mutations in the coding region, by changing the stop-codon read-through rate, or through co-option.

Attributing increases in evolvability to selection for evolvability is non-trivial. One method is to compare evolutionary outcomes from changing vs. static environments. If changing environments, where the ability to rapidly adapt is at a premium (Eshel 1973), give rise to enhanced evolvability, it seems plausible that selection for evolvability might be responsible. However, changing environments might have other, unintended side effects. Therefore, rigorous negative controls are needed for adaptive claims, including when using theoretical rather than empirical approaches (Wagner 2003; Artzy-Randrup *et al.* 2004; Solé and Valverde 2006; Lynch 2007a; Tsuda and Kawata 2010; Masel and Promislow 2016). To establish a negative control for selection for evolvability, we develop a model that incorporates a single polymorphism at a time through an origin-fixation approach (Gillespie 1982; McCandlish and Stoltzfus 2014). The adaptive evolution of evolvability requires some form of lineage selection, not necessarily involving groups, but requiring polymorphism in the evolvability allele to be simultaneous within a (meta-)population with polymorphism in the adaptations it helps promote. Origin-fixation simulations, in which such simultaneous polymorphisms are not allowed, are incapable of adaptive evolution of evolvability. Surprisingly, we show that enhanced evolvability via evolutionary capacitance can arise as a response to environmental change even when the adaptive evolution of evolvability is precluded.

## METHODS

### Genotype fitness

To capture the effects of recurrent environmental change on the evolution of cryptic sequences and stop-codon read-though rates, we build on the model of Rajon and Masel (2011). We focus on the evolution of the stop-codon read-through rate (*ρ*) and its effect on the expression of many cryptic sequences, each associated with one of *L_tot_* genes. To reflect observations that the distribution of fitness effects of new mutations is bimodal (Bloom *et al.* 2006; Eyre-Walker and Keightley 2007; Fudala and Korona 2009; Wylie and Shakhnovich 2011), we make alleles for each cryptic sequence either highly deleterious or largely benign. The fitness of an individual, *w*(*i*), is the product of four components. *w_m_* captures the damage from expressing deleterious cryptic sequences via read-through. *w_s_* encapsulates the cost of proofreading to avoid read-through. In order to prevent pathologically extreme cases of “error” rates greater than 0.5 from evolving, inherent costs of very high read-through rates are encapsulated by an additional term *w_h_*. Finally, we represent the fitness consequences of possessing a phenotypic trait value *x*, given the current environmental optimal value *o_e_* for that trait, through the term *w_e_*.

Stop-codon read-throughs occur with frequency *ρ*, and expose *L_del_* deleterious sequences out of the *L_tot_* cryptic loci. Deleterious sequences have an additive fitness cost *γ*=20 estimated from Geiler-Samerotte *et al.* (2011) and Xiong (2017), yielding:

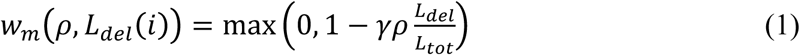

Next, we incorporate the cost of proofreading. Low error rates entail costly proofreading mechanisms that can slow reproduction (Fersht *et al.* 1982). In our model, additive investment in proofreading leads to a geometric decline in the error rate, making the cost of significant additional proofreading negligible at high read-through rates, but very costly as read-through rates approach zero. Using *δ* =10^−2.5^, a biologically reasonable decrease in growth rate of 0.72% ensues when read-through rates decrease from 10^−2^ to 10^−3^ (Rajon and Masel 2011):

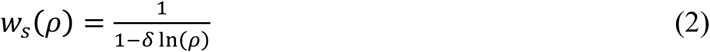

To ensure that “errors” do not become the new normal when all cryptic sequences are benign, i.e. to avoid *ρ* evolving values greater than 0.5 (Xiong *et al.* 2017), we use a phenomenological fitness term that has little impact on fitness for low values of *ρ*, but causes fitness to decline rapidly as the probability of a stop-codon read-through becomes large. The parameter *θ* determines the value of *ρ* at which fitness is zero; here we set *θ*=0.5. The remaining parameter, ***τ***,determines the rate at which fitness increases as *ρ* decreases. Here we use *τ*=20 to produce a function that is effectively neutral (i.e. *sN*≪ 1) when *ρ* <1/3 for population sizes between 10^3^ and 10^6^.

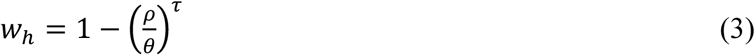

Finally, we incorporate the fitness effect of the match between a phenotypic trait x and the environmental optimum *o_e_*. Of the *L_tot_* loci, a subset of *K* loci have quantitative effects *α_k_* based on their regular coding sequences (Figure 2). Each of these *K* loci also makes a cryptic contribution *ρB_k_β_k_* to the phenotype. We use a binary *B* to represent the possibility that a trait-associated cryptic sequence can be highly deleterious in the same manner as any other among the *L_dei_* loci, and we use a real number *β* to represent the contribution of a cryptic sequence to the phenotypic trait. If highly deleterious, *B_k_* = 0 and the cryptic sequence does not contribute to the phenotype. If benign, *B_k_* = 1 and the cryptic sequence contributes *ρβ_k_* to the phenotype. Additionally, if a trait-associated cryptic sequence is highly deleterious, *B_k_* contributes to the overall cryptic sequence load *L_del_* in Equation 1. Fitness is a Gaussian function of the difference between the phenotype of each trait *x_i_* and the environmental optimum of that trait *o_e_*.

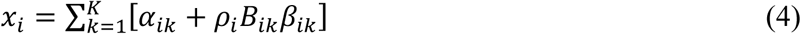

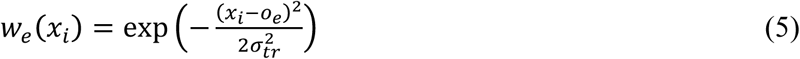

where *σ_tr_^2^* = 4. The overall fitness of genotype *i* is then a product of the four fitness components:

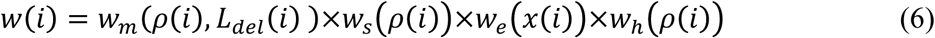

### Selection and drift

We used a single-deme origin-fixation (strong-selection weak-mutation) process (Gillespie 1982; McCandlish and Stoltzfus 2014). A mutation is generated, and then fixed or lost with probability determined by the Kimura equation (1962):

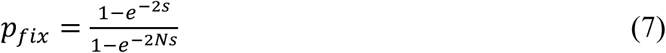

where *s* is the selection coefficient:

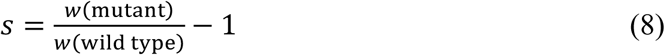

### Mutation

Our genotypes encode four evolving quantities: the stop-codon read-through rate (*ρ*), the number of loci that are fixed for a deleterious cryptic sequence (*L_del_*), the coding regions of trait-coding genes (*α*) and the cryptic regions of trait-coding genes (*β*). Following Rajon and Masel (2011), we assume that the cryptic sequences are on average 30 nucleotides long, making the rate of mutations in *L_tot_* cryptic loci *μ_Ldel_*=*L_tot_*×30×*μ* where *μ* is the mutation rate per base pair. Mutations in benign cryptic sequences convert the sequence to a deleterious state with probability *p_del_* = 0.4, and those in deleterious sequences convert the sequence to benign with probability *p_ben_* = 0.1. Therefore, in the absence of selection, a cryptic sequence has a probability *p_del_/(pd_el_+ p_ben_)*, or 0.8, of being deleterious. Assuming that the read-through rate is determined by relatively few genes (~3 with ~300 nucleotides each) yields a mutation rate in *ρ* of roughly *μ_p_*=1000*μ*. Mutations in the read-through rate change log_10_(*ρ*) by an amount sampled from a normal distribution with mean zero and variance 1. Assuming an average of 300 bp per locus and *K* total trait loci, mutations occur in non-cryptic sequences of trait-coding genes at rate *μ_α_*=300×*K*×*μ* and cryptic sequences at rate *μβ*=30×*K*×*μ*. A mutation in a trait-coding region has an additive effect according to a normal distribution with variance *σ^2^_tr_*=0.5/*K*. As in Rajon and Masel (2011), to prevent the unrealistic outcome of an unbounded increase in variance over time, we bias change in trait values toward zero. The mean effect of a mutation in a trait with coding value *α_k_* is thus -*α_k_*/50. Because we are using an origin-fixation process in which we allow for only one polymorphism at a time, the overall mutation rate *μ* has no effect.

### Environmental change

The environmental optimum changes every 1000 mutations by an amount sampled from a normal distribution biased to return the optimum toward zero. The rate of environmental change was chosen such that there are enough mutations to allow populations to adapt to one environmental change before the next occurs. Because we use an origin fixation process, the number of generations between mutations is not strictly defined. In a model with explicitly defined time, large populations generate more mutations per generation than small populations, and therefore should experience more mutations between environmental changes. By ignoring this effect, we effectively assume that enough mutations occur between environmental changes that any possibly fitness gains are saturated even in small populations.

To reflect findings that cryptic sequences seem to be maladaptive in new environments as often they are adaptive (True *et al.* 2004), the environmental optimum undergoes a random walk. The mean change in the environmental optimum, given the current environmental optimum *o_e_*, is -*o_e_*/5, producing a random walk biased toward zero. Simulations were run for 4×10^8^ mutations, i.e. 4×10^5^ environmental change events, which proved to be well in excess of convergence to a stationary distribution of states. Perl code used is available at github.com/pgnelson/emergent_capacitance.

## RESULTS

### Environmental change pushes the evolutionary dynamic toward the high-evolvability attractor

We simulated evolution using a single-deme, strong-selection weak-mutation model. When a mutation arises, it is either fixed or lost immediately, with outcome probabilities determined by the Kimura (1962) equation. Thus, our model does not allow for the standing genetic polymorphism in at least two loci, one the evolvability locus the other the adaptation it helps promote, required for the adaptive evolution of evolvability via lineage selection (see Figure 1). By only allowing a single mutation at a time, and using a single well-mixed population, we exclude both lineage selection and group selection and thus establish a rigorous negative control for the adaptive evolution of evolvability via cryptic sequences. Throughout this work, we use an organism’s robustness to stop-codon read-through, as measured by the proportion of benign cryptic sequences, as a proxy for the dramatic forms of evolvability documented in Rajon and Masel (2011). We focus on population sizes that represent the three regimes shown in Rajon and Masel (2011): low error rates (N=500), bistability (N=10^4^), and high error rates (N=10^5^).

We found that, despite our single-deme model being designed to preclude the adaptive evolution of evolvability, a rapidly changing environment caused evolution to the more evolvable, benign cryptic sequence, high error rate (Figure 3B) attractor in intermediate sized populations (N=10^4^), regardless of the initial conditions. Again, this result was obtained using a single-deme, strong-selection weak-mutation model, precluding selection for evolvability either between groups or between lineages as a cause. Because environmental change cannot directly affect selection on deleterious cryptic sequences, the root cause by which environmental change drives the evolution of high robustness and evolvability must derive from effects on the distribution of stop read-through rates.

**Figure 3:**
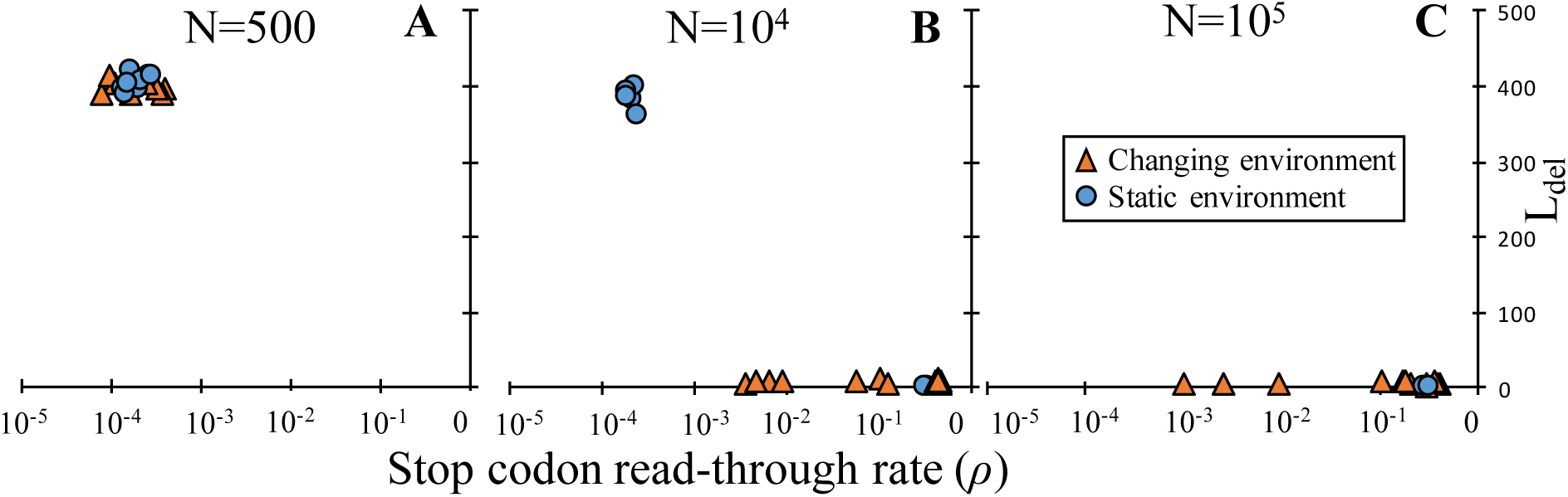
Environmental change facilitates the evolution of benign cryptic sequences and high error rates. In the absence of environmental change (circles), small populations (A, N=500) evolve low error rates coupled with mostly deleterious cryptic sequences, and large populations (C, N=10^5^) evolve mostly benign cryptic sequences coupled with relatively high read-through rates. When populations are of intermediate size (B, N=10^4^), outcomes can be bistable, evolving either mostly deleterious, or mostly benign cryptic sequences, depending on the starting conditions. A changing environment (triangles) has little effect on small populations (A) and leads to increased variance in the read-though rate in large population (C) but no change in the number of deleterious cryptic sequences. In intermediate-sized populations (B), a changing environment causes populations starting with mostly deleterious cryptic sequences to transition to a mostly benign, high-*ρ* state. Each circle or triangle represents the end value of an independent simulation run to equilibrium (1.5x10^8^ mutations).

Reading through a stop-codon can have two effects in our model. If a cryptic sequence is deleterious, a read-through event entails a fitness cost, independent of the environment. If a cryptic sequence is benign (*B_ik_*=1), expression can result in small, but potentially advantageous changes in the phenotype. Because we initialize our populations such that the phenotype imparted by the coding region of trait genes (∑*α_ik_*) is perfectly matched to the environment, expressing even benign cryptic sequences is unlikely to be advantageous in a static environment. The coding regions of trait genes may, however, become maladaptive when the environment changes, making expression of benign cryptic sequences potentially adaptive. Thus, if the net phenotypic effect of benign cryptic sequences (∑*B_ik_β_ik_*), or “net *β*”, moves an organism in the direction of a new environmental optimum (*o_e_*), selection may favor increased read-through rates despite the costs of expressing other deleterious cryptic sequences in the genome (*L_del_*).

### Environmental change leads to more extreme quantitative effects of cryptic sequences

In addition to purging deleterious cryptic sequences, environmental change has a marked effect on the evolution of benign cryptic sequences (*β*). In a static environment, where expression of cryptic sequences is never likely to be adaptive, net *β* is constrained around zero (Figure 4, left column). In contrast, a changing environment (Figure 4, right column) results in much more extreme values of net β. When the environment changes, the net effect of benign cryptic sequences is maladaptive roughly as often as it is adaptive. When maladaptive, selection maintains low read-through rates, and cryptic sequences (*β*) are primarily subject to drift and mutation, the latter weakly biased toward zero. When net *β* is adaptive, selection can favor high read-though rates, which can in turn expose cryptic sequences to selection. Furthermore, expressing cryptic sequences rarely brings a phenotype perfectly in line with a new environmental optimum, and mutations that, following environmental change, increase the overall magnitude of net *β* are often advantageous. Additionally, because environmental change undergoes a random walk where ∑*α_ik_* is too small in a new environment as often as it is too large, the sign of net *β* (positive or negative) does not affect its overall adaptive value. Thus, cryptic sequences tend to only be exposed to selection when extreme values are favored, producing a ratcheting effect leading to more extreme values, either positive or negative, under environmental change. Although via a different mechanism, our model concurs with the findings of Eshel and Matessi (1998) that a fluctuating environment leads to increased cryptic variation with a 50:50 chance of bias in either direction, creating a bimodal rather than normal probability distribution of possible states.

**Figure 4:**
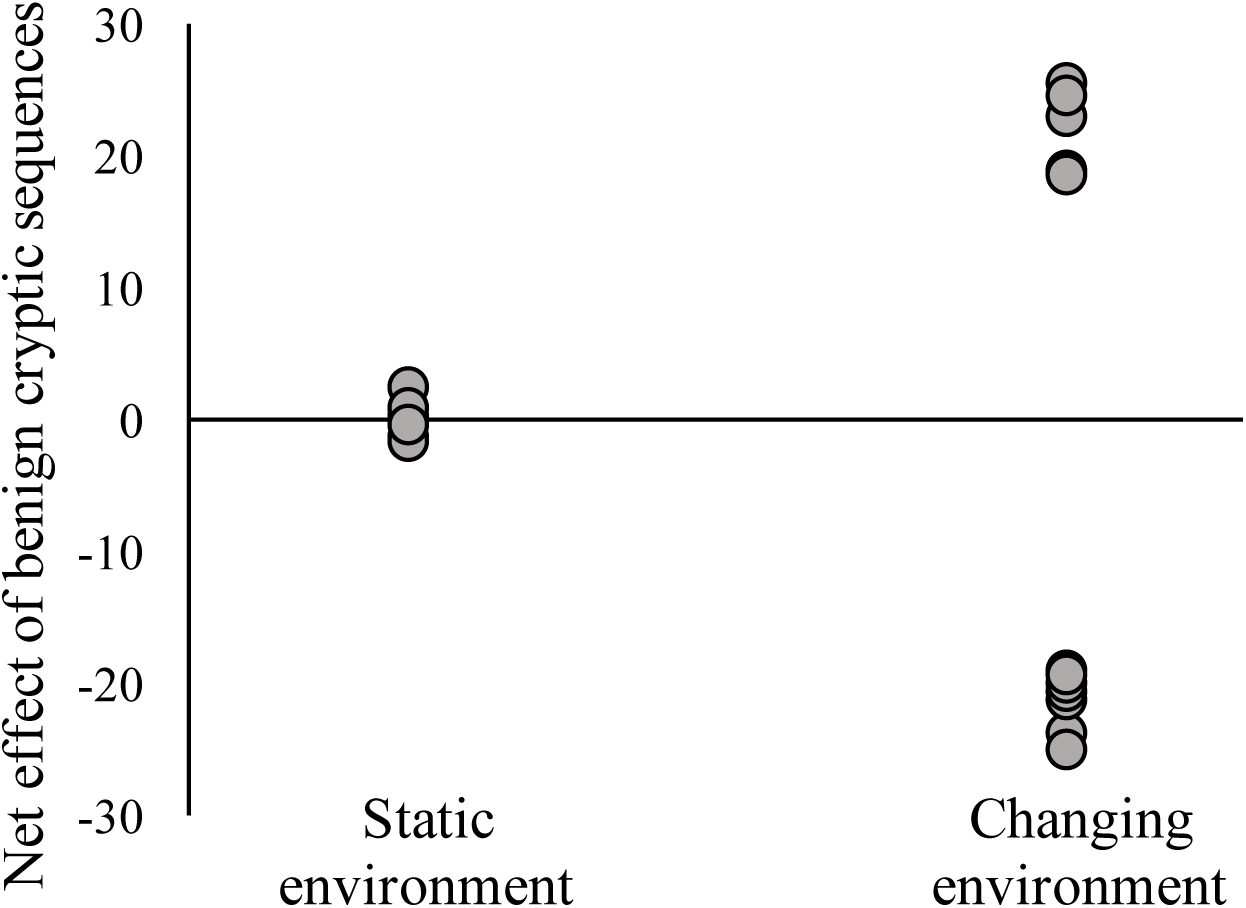
Changing environments result in more extreme cryptic sequences. Each circle shows the evolved net effect of benign cryptic sequences (∑*B_ik_β_ik_*) or net *β*, for an independent simulation of 1.5×10^8^ mutation trials where *N*=10^4^.

### Dissecting the effect of environmental change on the evolution of stop-codon read-through rates

To investigate precisely how environmental change affects the distribution of stop-codon read-through rates, the simulation was re-run with the number of deleterious cryptic sequences fixed, which we refer to below as a “fixed *L_del_* simulation”. We ran three independent fixed *L_del_* simulations for a sample of *L_del_* values between zero and the equilibrium expected by mutation bias (*L_del_* = 400, 375, 350…0). By preventing the evolution of deleterious cryptic sequences, we isolated the effect of environmental change on read-through rates (Fig. 5 A-B). For each simulation, we recorded the overall distribution of *ρ* (Fig. 6), the frequency of transitioning to a high *ρ* state (*ρ* > 10^−2.5^) and a time series of *ρ* while in a high *ρ* state.

**Figure 5:**
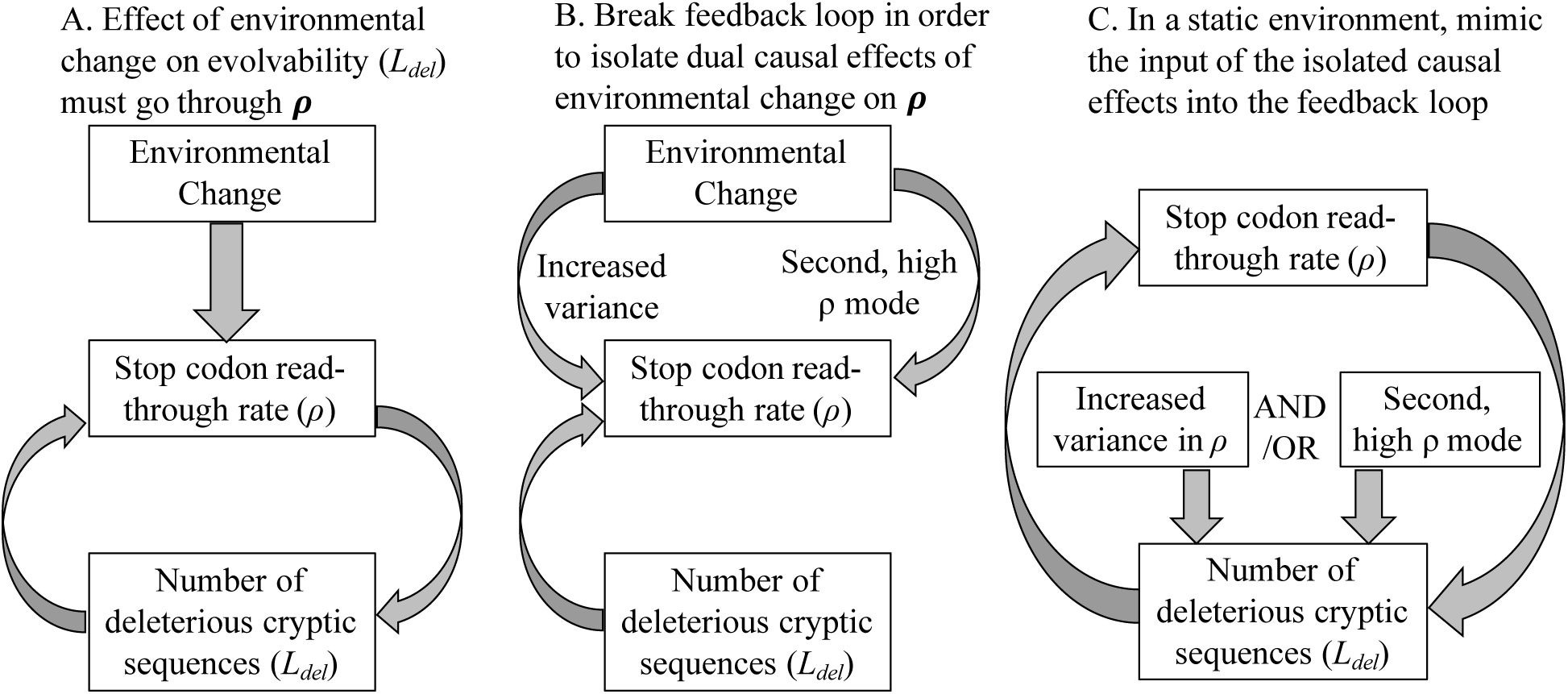
To determine if the evolution toward the benign cryptic sequence attractor is caused by the increase in the variance of *ρ*, or by the appearance of the second, high *ρ* mode, we dissect the effects of recurrent environmental change on the feedback loop between *ρ* and *L_del_*. A) Environmental change cannot directly change selection on *L_del_*, only indirectly via changes in *ρ*. B) We determined the direct effect of environmental change on the stop-codon read-through rate in a system constrained to prevent the evolution of deleterious cryptic sequences. To achieve this constraint, we set the mutation rates between benign and deleterious cryptic sequence states (*L_del_*) to zero and recorded the distribution of *ρ* at steady state. C) Instead of letting environmental change drive change, we mimicked distinct aspects of its effects on *ρ*, to simulate the evolution of deleterious cryptic sequences in the presence of these drivers. This enabled us to determine which aspect is responsible for the shift to benign cryptic sequences under environmental change.

**Figure 6:**
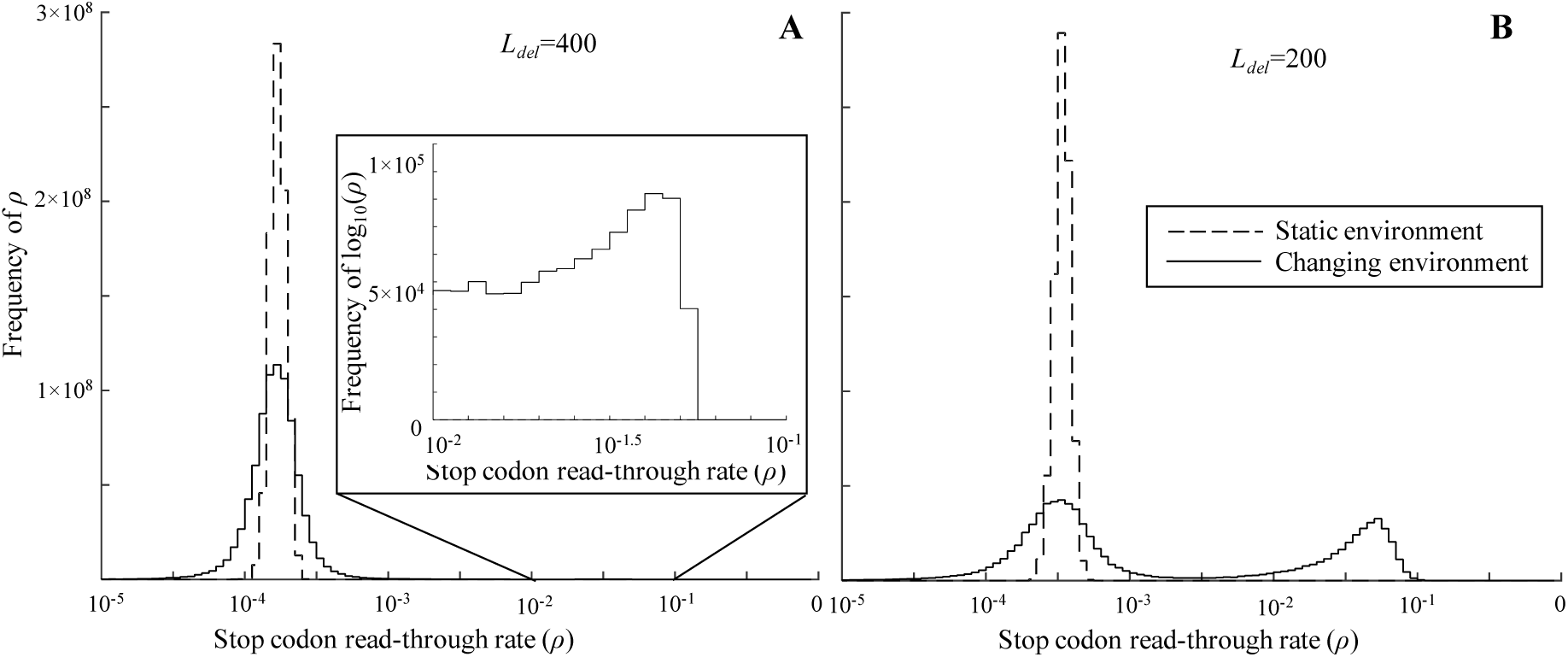
Effect of environmental change on the distribution of stop-codon read-through rates (*ρ*) for two values of *L_del_* (400, 200). To isolate the effect of environmental change on stop-codon read-through rates, the number of deleterious cryptic sequences was held constant. The dashed line shows that the read-through rate evolves to the low attractor with relatively low variance. Environmental change (solid) yields a distribution around the low *ρ* attractor with similar mean but higher variance. The inset in panel A shows the emergence of a second, low frequency mode at high read-through rates (*ρ*~l0^−1.4^) under environmental change. The frequency of this high *ρ* mode increases markedly when the read-though rate is less constrained by deleterious cryptic sequences (Panel B, *L_del_* = 200).

In a static environment (Figure 6, dashed lines) the stop-codon read-through rate is unimodal around the low error rate attractor (*ρ*~10^−3.8^ for *L_del_* = 400; *ρ*~10^−3.5^ for *L_del_* = 200). Environmental change has two effects on the distribution of read-through rates (Figure 6, solid lines). First, environmental change causes the variance of log_10_(*ρ*) around the low error rate attractor to increase ten-fold for *L_del_* = 400 and 100-fold for *L_del_* = 200. Second, a smaller, second mode arises at high read-through rates (*ρ*~10^−1.2^), the frequency of which increases as *L_del_* decreases.

Furthermore, by examining the timecourse of *ρ*, we see that the error rate under environmental change has the hallmarks of an evolutionary capacitor (Masel 2013), with high error rates being clustered in episodes, or bouts. When the environment changes such that expressing cryptic sequences becomes on balance advantageous, selection can favor dramatically increased read-through rates (Figure 7, solid line). When environmental change creates a large mismatch between the trait coding values and the environmental optimum (Figure 7, dashed line), high read-through rates can increase rapidly to tap into benign cryptic sequences of trait coding genes (net *β*). However, increased read-through rates come with the costs of expressing deleterious cryptic sequences. Therefore, as adaptations in trait-coding sequences (*α*) accumulate, read-through rates decrease and eventually return to the low *ρ* attractor.

**Figure 7:**
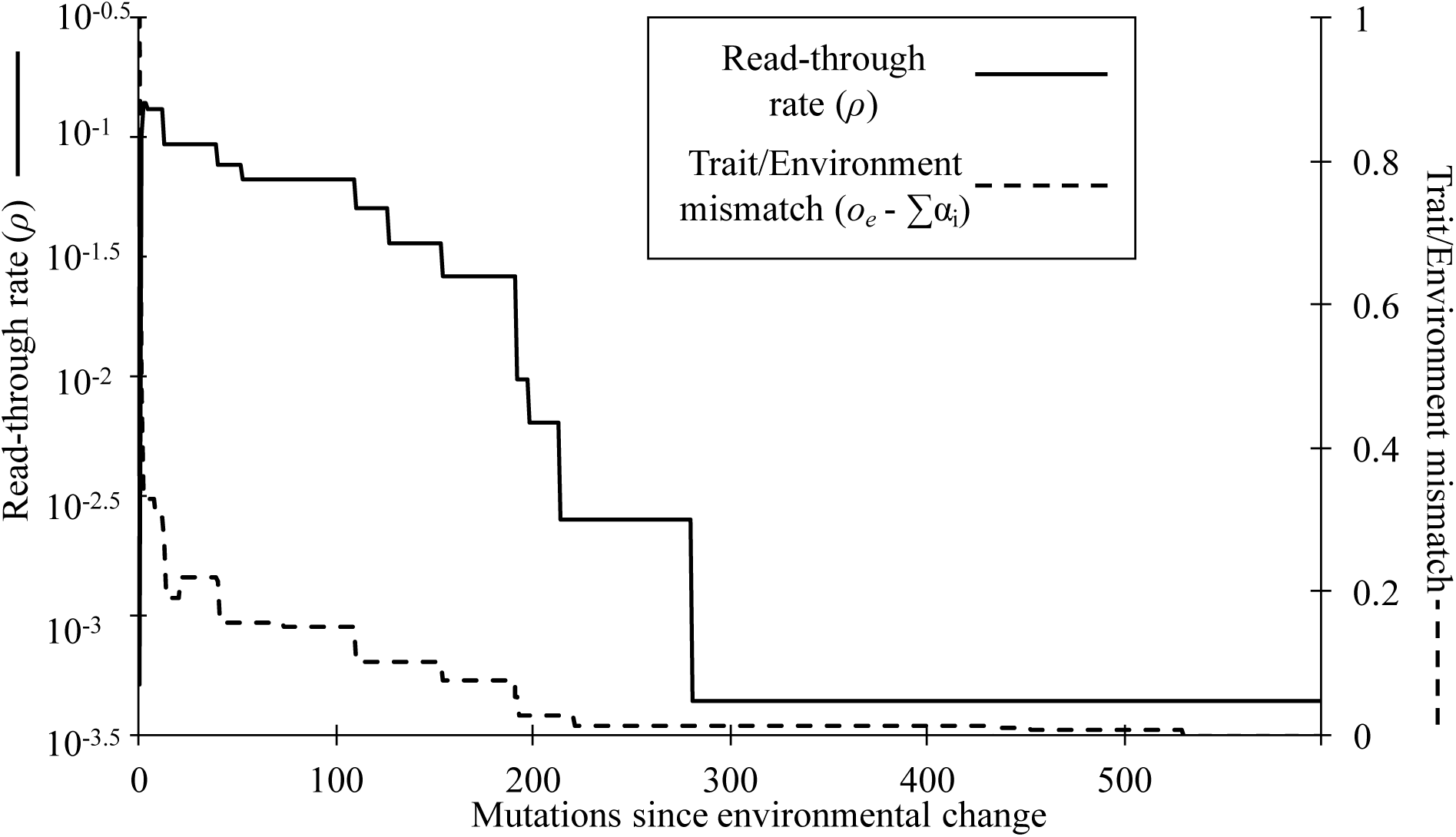
Transient elevation of stop-codon read-through rates acts as an evolutionary capacitor under environmental change. The stop-codon read-through rate (*ρ*, solid line) increases immediately after an environmental change (0 on the x-axis). As the mismatch between the environmental optimum and the trait values of the coding region ((*o_e_* - ∑α_i_), dashed line) decreases, *ρ* decreases, returning to the low error attractor. This representative timecourse illustrates one bout of high read-through rates in a simulation with *L_del_* = 400 under environmental change. Note that in our simulation, the environment changes every 15,000 mutations, making the duration of this capacitance bout relatively short compared to the total consecutive time spent in a single selective environment.

Environmental change therefore begets transient runs of high stop-codon read-through rates during which selection may efficiently purge deleterious cryptic sequences. Next, we ask whether it is the rare bouts of high read-through rates, or the increase in variance around the low *ρ* mode, that is the causative factor in the evolution of more benign cryptic sequences.

### Rare bouts of high read-through rates drive the transition

To determine which characteristic(s) of the environmental-change-driven *ρ*-distribution are causally responsible for the evolution of more benign cryptic sequences, we tested our model under four different manipulations of *ρ* (listed below), each based on the fixed *L_del_* simulations outlined above. Based on the observed frequency of *ρ* values under environmental change (Figure 6), we used a cutoff of *ρ* =10^−2.5^ to separate the low *ρ* distribution from the emergent high *ρ* distribution. To track the coevolution of *ρ* and *L_del_*, our manipulations are based on values of *ρ* taken from the fixed *L_del_* simulations with the closest number of deleterious cryptic sequences to the evolved value of *L_del_*. For example, when evolved *L_del_*=390, we use values of *ρ* drawn from simulations with fixed *L_del_*=400.

We found that the bouts of high read-through rates engendered by environmental change are both necessary and sufficient to purge deleterious cryptic sequences (Figure 8). Neither increased variance around the low read-through rate attractor, nor evenly distributed occurrences of high *ρ* values, nor a combination of both result in a transition to mostly benign cryptic sequences. These results show that the episodes of increased rates of stop-codon read-though caused by environmental change act as an evolutionary capacitor, allowing rapid adaptation to a new environment. As a side effect, they purge deleterious cryptic sequences, pushing the evolutionary dynamic toward the high error rate, high evolvability attractor.

**Figure 8:**
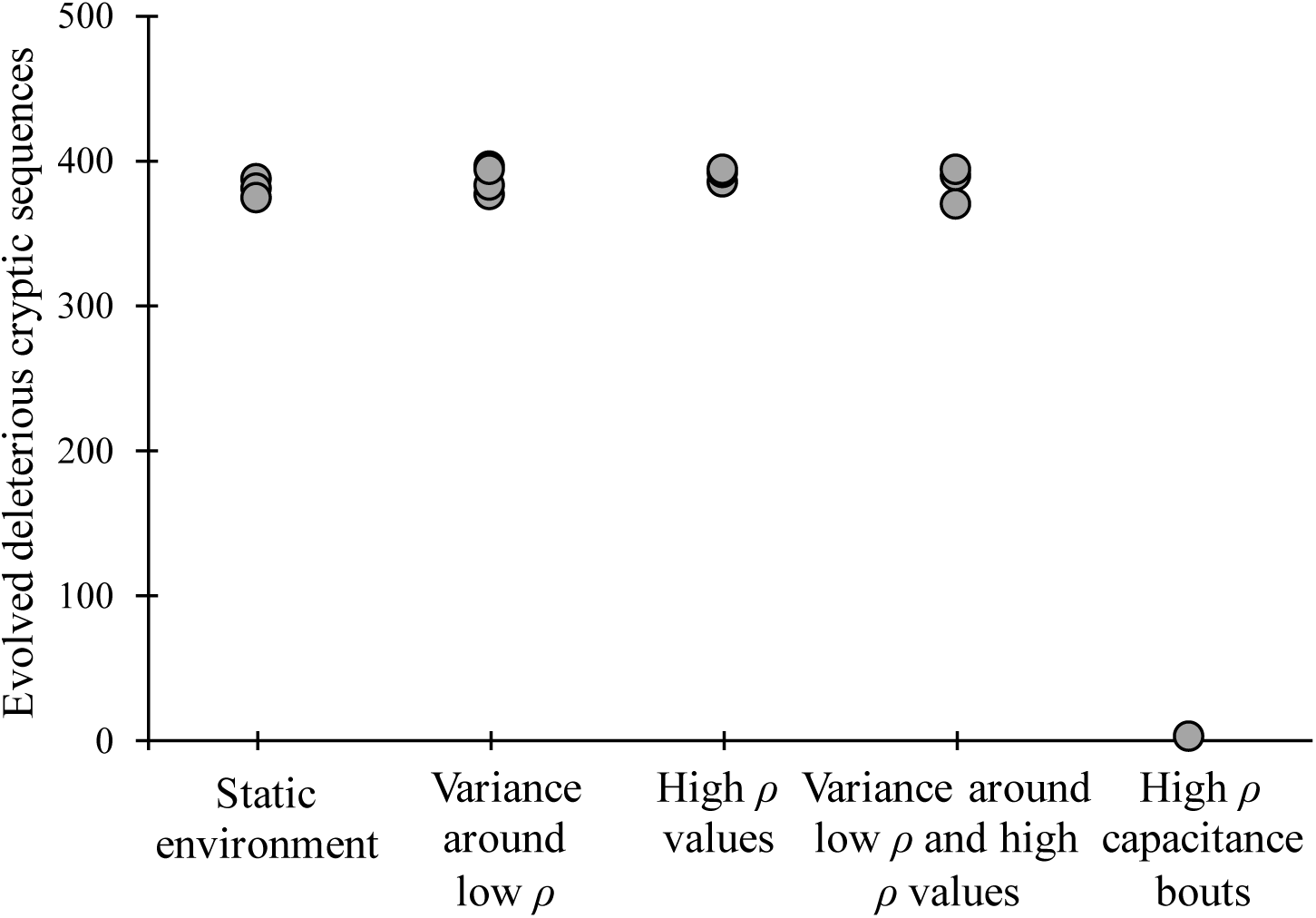
Effect of rare occurrences of high stop-codon read-through rates on the evolution of cryptic sequences. We ran our model under four conditions, with evolution under a static environment for comparison, using *N*=10^4^ to simulate isolate the various effects of environmental change. First, we simulate the effect of increased variance around the low error rate attractor by adding to the log_10_ of the read-though rate *ρ* a random number drawn from a normal distribution, with mean 0 and variance equal to the increase in variance in *ρ* due to environmental change. Second, we sample from the high error rate mode that emerges under environmental change. Read-through rates were sampled from the high *ρ* distribution at the frequency of *ρ* >10^−2.5^ observed in the fixed *L_dei_* simulations, otherwise un-manipulated, evolved values of *ρ* were used. Third, we test the effect of the high *ρ* mode together with increased variance on the evolution of deleterious cryptic sequences. Finally, we test the effect of high *ρ* capacitance bouts by injecting time series of high *ρ* values. Every 15000 mutations, populations may enter a high *ρ* state with the same probability as observed in the fixed *L_del_* simulations. When a population enters a high *ρ* state, a series of high *ρ* values is drawn from those observed in the fixed *L_del_* simulations. When the series of high *ρ* values is exhausted, the un-manipulated, evolved value of *ρ* is used again. In all four cases, we manipulated the value of *ρ* used for the purposes of determining the probability of fixation of mutations to cryptic sequences. Each marker indicates the evolved number of deleterious cryptic sequences after 1.5×10^8^ mutations.

## DISCUSSION

Our finding that a changing environment engenders bouts of very high read-through rates is similar to the phenomenon known as evolutionary capacitance (Masel 2013). Capacitance is the switching on and off of the phenotypic expression of cryptic variants (Masel 2013), generally in stressful environments (Sangster *et al.* 2004), due to a capacitor, meaning a widget that can toggle cryptic expression. Evolutionary capacitance can occur in sexual organisms, via recombination-driven variation (Sangster *et al.* 2004; Masel 2006; Schlichting 2008; Masel and Trotter 2010), and in asexual organisms, through mutation-driven variation (Griswold and Masel 2009; Masel and Trotter 2010). Evolutionary capacitors enhance evolvability by maintaining phenotypic fidelity under benign conditions, while increasing variation when conditions become difficult (Tyedmers *et al.* 2008), or indeed occasionally at random as a bet-hedging strategy against environmental change (Masel 2005; King and Masel 2007).

We found that in a changing environment, capacitance-like behavior (fluctuating error rates that purge deleterious cryptic variation) emerges without the benefit of a widget to toggle robustness. Emergent capacitance was observed in populations with drift barrier effective population sizes of roughly one order of magnitude around *N_e_* =10^4^, which encompasses estimates for effective population sizes in humans and other vertebrates (Sung *et al.* 2012). The scope of these findings depends on the fitness costs of deleterious cryptic sequences. Highly deleterious cryptic sequences are readily purged, where more mildly deleterious cryptic sequences require higher effective population sizes (Xiong *et al.* 2017). Therefore, the effective population size at which emergent capacitance may arise likely varies with the distribution of fitness effects of cryptic sequences.

Evolutionary capacitors enhance evolvability by tapping into stores of benign cryptic variation. When in a cryptic state, genetic variation can accumulate due to drift, either among organisms in the case of sexual organisms, or among alternative cryptic sites in a genome in the case of asexual organisms. Because most new variation is likely to be deleterious, co-option / genetic assimilation, i.e. constitutive expression of a cryptic variant, is likely inviable (Figure 9, middle column). When robustness is temporarily relaxed, however, deleterious cryptic sequences can be purged by selection, leaving behind only benign cryptic variants (Figure 9, right column) (Rajon and Masel 2011). Thus, capacitance enhances a species’ adaptive potential in two ways. First, capacitance can engender rapid (Rando and Verstrepen 2007), but temporary (Masel and Bergman 2003), change in phenotype under stressful conditions (True and Lindquist 2000; True *et al.* 2004; Tyedmers *et al.* 2008). Second, benign cryptic phenotypes may be permanently coopted by mutation (Griswold and Masel 2009; Masel and Trotter 2010) or recombination.

**Figure 9:**
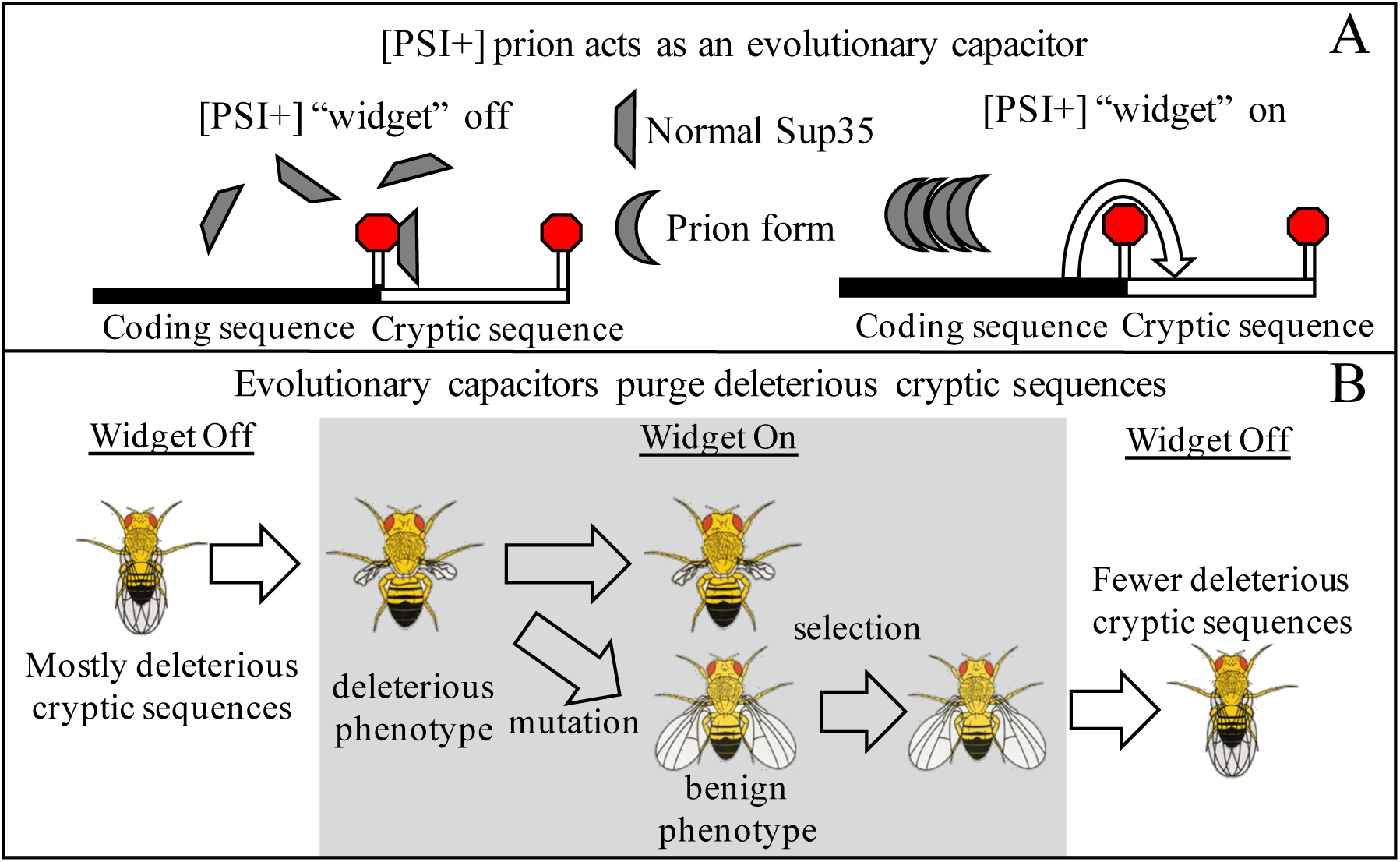
The [PSI+] prion acts as an evolutionary capacitor, leading to more benign cryptic sequences. In its normal conformation, Sup35 facilitates translation termination at stop codons. In its alternate form as the [PSI+] prion, Sup35 is sequestered and stop codon read-through rates are higher (A). When a capacitance widget such as [PSI+] decreases robustness, cryptic phenotypes are expressed, allowing selection to purge deleterious cryptic sequences in favor of benign cryptic sequences (B). Images modified from themadlolscientist (2008).

Capacitance hinges on a breakdown in robustness exposing cryptic variants to selection, and then a restoration of robustness making most variants cryptic again, except for those that have been co-opted through mutation or genetically assimilated through recombination. Therefore, a key aspect of capacitance is that it must be possible to switch robustness on and off in a reversible fashion (Lancaster and Masel 2009). For example, while robustness can easily be switched off by knocking out genes within complex regulatory systems, potentially leading to capacitance-like phenomena (Bergman and Siegal 2003), gene knockouts are not easily reversible. Because there is no mechanism to return knockout-revealed variants to a cryptic state, knockout-induced capacitance is unlikely to lead to enhanced evolvability. Therefore, a defining feature of evolutionary capacitance is a capacitor, a widget capable of mediating the switch between cryptic and revealed states (Ruden *et al.* 2003). The best-established candidate for adaptive capacitance is the yeast prion [PSI+], precisely because the epigenetic mode of prion inheritance allows robustness to be reversibly switched both off and on. The presence of [PSI+] increases the rate of stop-codon read-through, reducing robustness to the consequences of sequences lying beyond stop-codons. While it has been shown mathematically that capacitance can (Masel 2005; King and Masel 2007) and in the case of [PSI+] likely did (Masel and Bergman 2003) arise due to selection for enhanced evolvability, a non-adaptive exploration of the evolution of capacitance via increased molecular error rates has not yet been conducted.

Due to our weak-mutation strong-selection assumption, which naturally precludes linkage between loci and therefore the ability of recombination to explore new phenotypes, our findings are most applicable to less sexual systems, chiefly the [PSI+] prion in yeast. The [PSI+] prion is an aggregated form of the Sup35 protein, with an alternate conformation (Wickner *et al.* 1995). In its normal form, Sup35 facilitates stop-codon termination in yeast (Stansfield *et al.* 1995; Zhouravleva *et al.* 1995). Because Sup35 cannot perform its normal function while in an aggregate, the [PSI+] prion results in a dramatic increase in stop-codon read-through rates (Firoozan *et al.* 1991) (Figure 9A). Such high read-through rates can occasionally be adaptive for certain environments and in certain genetic backgrounds (True and Lindquist 2000; Joseph and Kirkpatrick 2008). Furthermore, the [PSI+] prion preferentially appears in stressful environments (Tyedmers *et al.* 2008; Doronina *et al.* 2015). Thus, [PSI+] is known as a phenotypic or evolutionary capacitor (Masel 2013) because it acts in a concerted way to switch cryptic sequences at many loci simultaneously from a high-latency (low *ρ,β* is neutral) to low-latency (high *ρ,β* is phenotypically meaningful) state. If the net effect is adaptive, this buys the lineage time as a stop-gap, accelerating eventual adaptation via co-option. Given the existence of benign cryptic sequences, which themselves increase evolvability, the existence of a capacitor as a widget to exploit them increases evolvability further (Masel and Bergman 2003; Masel 2005; Griswold 2006; Masel 2006; King and Masel 2007).

Although we did not construct our current model with the intent of mimicking the behavior of the [PSI+] prion, the effect of environmental change on the stop-codon read-through rate bears a striking resemblance to the dynamics of the [PSI+] prion in yeast. Our model found, unexpectedly, that environmental change results in a bimodal distribution of read-through rates, either very high or very low, similar to the dichotomy caused by the presence vs. absence of [PSI+]. Thus, our results suggest that the [PSI+] prion may be filling a niche created by a changing environment.

In our simulations, we made the permissive assumption of mutational symmetry in the evolution of *ρ*. In reality, it is much harder to lower error rates than to raise them. Adding a mutation bias toward higher error rates would cause organisms to spend longer in the high error rate state, and thus more rapidly purge deleterious cryptic sequences. Conversely, if a widget allows an organism to rapidly return to a low error rate state, deleterious cryptic sequences may be able to persist despite evolutionary capacitance. Note that previous work has shown that a reversible epigenetic switch like [PSI+] evolves more easily when the alternative path to adaptation involves relatively irreversible loss-of-function mutations (i.e. those favored by strong mutation bias) (Lancaster and Masel 2009).

Our origin fixation model necessarily precludes recombination, making our results most directly comparable to species that undergo relatively little sex (Griswold and Masel 2009; Tsai *et al.* 2012). However, evolutionary capacitors are certainly not restricted to asexual species. Molecular chaperones such as Hsp90 (Sangster *et al.* 2004; Chen and Wagner 2012; Siegal and Masel 2012) have been purported to act as capacitors in sexual species by buffering the effect of mutations, and then exposing phenotypic variation under stressful conditions or when disrupted through laboratory manipulation. Loss of function driven capacitance is not limited to chaperones and may be a feature of many genes (Levy and Siegal 2008; Takahashi 2013; Taylor *et al.* 2016). Theory suggests that capacitance may be a general feature of gene networks under selection for robustness to developmental noise (Bergman and Siegal 2003). Further research is needed to determine whether the particular phenomenon that we document here, emergent capacitance, might also occur in populations with high rates of recombination. Whereas asexual populations rely on mutation to produce new genetic variation, in sexual populations the genetic variance produced by new mutations may be dwarfed by the variance generated by recombination (Masel and Trotter 2010). Therefore, explicit modeling of recombination, coupled with a careful, negative control for the adaptive evolution of evolvability, is necessary to determine if the phenomenon of emergent capacitance due to environmental change is only relevant to asexual organisms, or whether it is a universal phenomenon.

## Acknowledgements

We thank Kun Xiong for helpful exchange of methods and the John Templeton Foundation (39667) and the PERT program (NIH K12 GM000708) for funding.

